# Chromosome-level Genome Assembly and Annotation of the Degu (*Octodon degus*)

**DOI:** 10.64898/2026.01.22.701162

**Authors:** Jeffrey H. Chiu, Nathan R. Zemke, B. Maximiliano Garduño, Bing Yang, Melanie L Oakes, Weronika Bartosik, Kristie Wong, Patricio Pezo-Valderrama, Yang Xie, Claudio Urra, Juan Macias, Chad Tomlinson, Patricia Cogram, Xiangmin Xu, Bing Ren

**Affiliations:** Department of Cellular and Molecular Medicine, UC San Diego, School of Medicine, La Jolla, CA USA; Biomedical Sciences Graduate Program, University of California, San Diego, La Jolla, CA; Center for Epigenomics, University of California San Diego, La Jolla, CA, USA; Department of Anatomy and Neurobiology, University of California, Irvine School of Medicine, Irvine, CA, USA; Department of Biological Chemistry, University of California Irvine, Irvine, CA 92697; Institut de Biologia Evolutiva (CSIC/Universitat Pompeu Fabra), Barcelona, España; New York Genome Center, New York, NY, USA; IEB, Faculty of Science, University of Chile, Santiago, Chile; McDonnell Genome Institute, Washington University School of Medicine, St. Louis, MO 63108, USA; Department of Genetics, Washington University School of Medicine, St. Louis, MO 63108, USA; Scientific and Technological Center, Fundación Ciencia & Vida, Santiago, Chile; The Center for Neural Circuit Mapping, University of California, Irvine, CA, USA; Department of Genetics and Development, Systems Biology, Biochemistry and Molecular Biophysics, Columbia University Irving Medical Center, New York, NY, USA

## Abstract

High-quality reference genomes are essential for comparative and functional genomics yet remain unavailable for many emerging model organisms. Here, we report OctDeg2.0, a chromosome-level genome assembly and annotation of the degu (*Octodon* degus), generated using PacBio HiFi long-read sequencing and Hi-C scaffolding from a single male individual. *Octodon degus* is of growing interest as a natural animal model for aging and Alzheimer’s disease research. The 3.4 billion base-pair assembly comprises 28 autosomes and both sex chromosomes, with markedly improved contiguity and completeness over the prior short-read assembly while maintaining high base accuracy. The new assembly provides substantially enhanced gene annotation, clearer resolution of chromatin architecture and cis-regulatory element landscapes, with much improved characterization of repetitive and structurally complex regions, including centromeres and segmental duplications. We present a streamlined and reproducible pipeline for high-resolution *de novo* genome assembly and gene annotation applicable to other emerging model organisms. This work establishes a robust genomic resource for studying molecular and cellular mechanisms in health and disease in *Octodon degus*.

## Introduction

The common degu (*Octodon degus*) is a medium-sized (170-300 grams) rodent endemic to the Mediterranean savanna of central Chile. First described in 1782 by the Chilean-Spanish naturalist Abate Molina, the *Octodon degus* belongs to the family Octodontidae, which comprises 14 species (Opazo, 2005; Woods & Boraker, 1975). Degus, along with guinea pigs and chinchillas, are included within the New World parvorder Caviomorph. Together with the Old World parvorder Phiomorpha, which includes the naked mole rat, they form the infraorder Hystricognathi (Opazo, 2005). These herbivores live in self-built burrows and are active during the day, feeding on a variety of shrub foliage, seeds, and connective stem tissues (Meserve et al., 1983).

Degus are highly social, intelligent, and long-lived rodents. They typically live in colonies of three to seven individuals and communicate using a diverse repertoire of 15 vocalizations, including squeals, barks, and groans (Long, 2007). Their high cognitive abilities have been demonstrated in various memory and behavioral tests (Garduño et al., 2025). Notably, degus were the first rodents shown to use tools and understand their physical properties after extensive training (Okanoya et al., 2008). In captivity, they typically live up to five to eight years but have been reported to reach 14 years of age (Garduño et al., 2025). These characteristics highlight the common degu as a valuable model organism for studying cognition and aging.

Aged degus have been found to exhibit neurological and histological changes mirroring those seen in Alzheimer’s disease (AD) (Garduño et al., 2025). A 2012 longitudinal study on degus aged 6 to 60 months found three general age-associated trends: cognitive decline in spatial memory and object recognition memory, increased impairments in CA3-CA1 synaptic transmission and plasticity, as well as elevated levels of soluble Aβ oligomers and tau phosphorylation (Rivera et al., 2021). Recent studies have also found that wild-type aged degus (4 to 5.5 years) exhibit a wide range of burrowing performance, where animals unable to burrow more than 25% of the pellets in a burrowing tube have significantly increased levels of Aβ deposition and Tau pathology (Tan et al., 2022). An additional study of genomic variations in this outbred degu population identified novel *Apoe* variants associated with increased Aβ plaques (Hurley et al., 2022). These findings provide a strong argument for the degu as a promising natural animal model for studying sporadic Alzheimer’s disease. To realize its potential as an AD animal model, an accurate and contiguous *O. degus* genome is needed.

The previously available degu genome assembly, OctDeg1.0 (NCBI RefSeq: GCF_000260255.1), was released by the Broad Institute in 2012. This assembly was generated from approximately 80x coverage Illumina HiSeq short-read data using the ALLPATHS assembly pipeline as part of a whole-genome shotgun sequencing effort that was not formally published. Although a significant resource for the field, OctDeg1.0 is highly fragmented (>7000 scaffolds), limiting comprehensive analyses of repetitive genomic regions and higher-order chromosomal organization. Here, we present a detailed, straightforward genome building pipeline used to construct the first chromosome-level *Octodon degus* genome (Supplemental Fig. 1), which we will refer to as OctDeg2.0.

This new primary assembly, OctDeg2.0, has a haploid number of 29 (28 autosomes + 1 sex chromosome) and a haploid genome size of ∼3.4 billion base pairs (Gbp). It was generated using ∼45x coverage of PacBio HiFi long reads and ∼100x coverage of Hi-C short reads from a single male degu. Compared to short-read-based OctDeg1.0, OctDeg2.0 exhibits substantially improved contiguity and completeness. This enhanced assembly not only improves gene annotation and the interpretation of epigenetic signals but also enables robust analysis of highly repetitive genomic regions, including centromeres and segmental duplications. We anticipate that this chromosome-level genome will serve as a valuable resource for the scientific community, facilitating the identification of the genetic and epigenetic basis underpinning age-associated histological and cognitive heterogeneity of *Octodon degus*.

## Result

### Chromosome-level genome assembly

In Fig. 1A, we detailed the pipeline used to build the chromosome-level genome assembly, which includes four major steps: 1) assembly, 2) mitochondrial contig filtering, 3) scaffolding, 4) sex chromosome identification. This approach is adapted from established practices used in recently published vertebrate genome assemblies (Lok et al., 2017; Okuno et al., 2023; Sokolowski et al., 2024).

**Figure 1.**
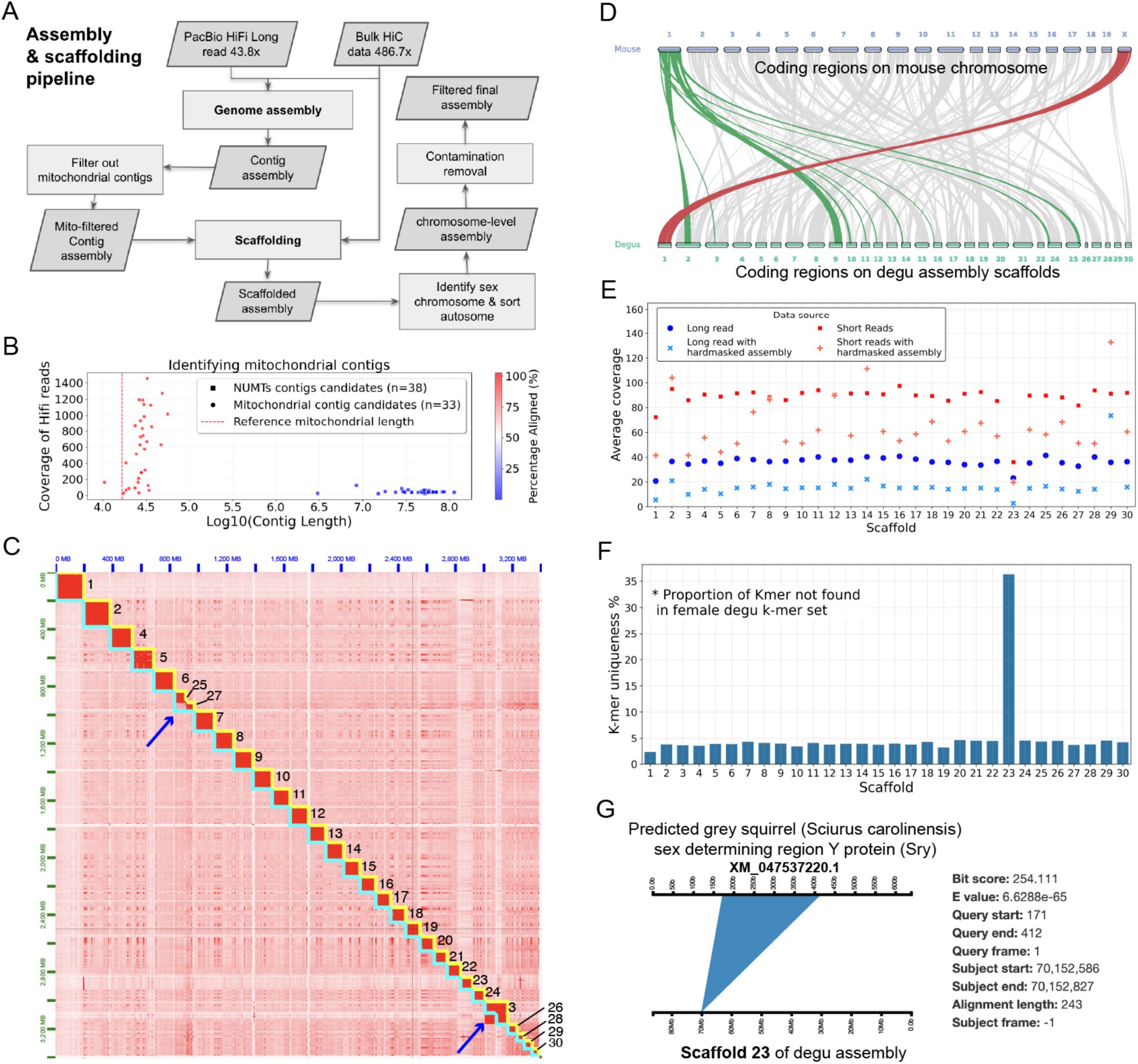
Chromosome-level genome construction using *de novo* genome assembly pipeline. A) Assembly and scaffolding pipeline using 43.8x PacBio HiFi long reads and 486.7x Hi-C short reads as input. Specifically ∼450x Hi-C were used for assembly and ∼60x Hi-C were used for scaffolding. B) Scatterplot of the mitochondria-associated contigs from which contigs with high similarity and length close to the mitochondria genome were filtered out. The red dash line is the length of the known degu mitochondria genome. C) Hi-C matrix showing the chromosomal blocks after HapHiC scaffolding. Blue arrows indicate manual curation sites to adjust the position of the scaffold boundaries from the cyan chromosome boundaries to gold chromosome boundaries. The numbers indicate the relative size of each scaffold from largest (1) to smallest (30). D) Synteny plot showing exclusive synteny between the X chromosome of the mouse genome and scaffold 1 of the degu assembly. E) Dotplot showing the respective read coverage of each chromosome for long read (dark blue), long read with hardmasked genome (light blue), short read (red), and short read with hardmasked genome (orange). F) Barplot of the proportion of k-mers not found in the k-mer set derived from the female genome. G) Blast result of grey squirrel *Sry* gene alignment to scaffold 23 of the new assembly.

Using HiFi long reads and Hi-C short reads as input, we assembled a 3.4 Gbp *Octodon degus* genome with a GC content of 41.34%. Our new contig-level assembly of the *Octodon degus* represents a major improvement over the existing scaffold-level OctDeg1.0 reference, increasing the contig N50 by more than 4-fold (from 12 Mbp to 50 Mbp) and reducing the number of sequences by over 10-fold (from 7,135 to 524) (Table 1). The new assembly also has substantial improvement in completeness and quality. Analysis using BUSCO (Manni et al., 2021) reveals an increase in complete markers from 97.8% to 98.6%, and decreases in fragmented (-0.5%), missing (-0.3%), and erroneous (-1.2%) BUSCO genes (Table 1). Evaluating the genome using Illumina short reads reveals a 5% increase in k-mer completeness and an improvement in basepair accuracy, with a Phred score (QV) increase from 31.5 to 32.1.

**Table 1.** Assembly metrics for OctDeg1.0 and the new *Octodon degus* genome assembly at contig- and scaffold-level resolution. Metrics are shown for the original OctDeg1.0 scaffold-level assembly and for contig- and scaffold-level assemblies generated in this study. Scaffold-level statistics for the new assembly exclude mitochondrial contigs and reflect the pre– contamination-filtered assembly. The final contamination-filtered assembly has a total length of 3,400,334,070 bp and comprises 200 sequences. Assembly quality was assessed using BUSCO and Merqury.

|  | OctDeg1.0 | Assembly from the current study |  |
| --- | --- | --- | --- |
|  | Scaffold | Contigs | Scaffolds (excluding mitochondrial contigs) |
| Total length (bp) | 2,995,889,303 | 3,404,567,379 | 3,403,620,422 |
| Number of sequence | 7135 | 524 | 288 |
| Largest sequence | 41,204,401 | 112,645,956 | 197,914,338 |
| N50 | 12,091,372 | 50,524,101 | 125,803,812 |
| L50 | 77 | 26 | 12 |
| GC content | 41.5 | 41.33 | 41.34 |
| Number of sequences > 10 Mb | 110 | 72 | 30 |
| Number of sequences > 25 Mb | 17 | 53 | 30 |
| % sum length of sequences > 10 Mb | 62.40% | 89.37% | 99.35% |
| % sum length of sequences > 25 Mb | 17.99% | 80.34% | 99.35% |
| Busco complete | 97.8% | 98.60% | 98.70% |
| Busco complete & single copy | 92.9% | 90.0% | 90.10% |
| Busco complete & duplicated copy | 4.9% | 8.60% | 8.60% |
| Busco fragmented | 1.2% | 0.70% | 0.7% |
| Busco missing | 1% | 0.70% | 0.6% |
| Busco erroneous | 5.70% | 4.50% | 4.50% |
| Merqury kmer completeness | 82.85% | 87.72% | 87.72% |
| Merqury QV estimation | 31.5 | 32.1 | 32.1 |
| Merqury error rate estimation | 0.0709% | 0.0610% | 0.0610% |

To prepare the contig assembly for genome scaffolding, we identified and filtered out 33 high-likelihood mitochondrial contigs (Fig. 1B). Using Hi-C reads, which includes additional Hi-C reads not used in the assembly, we scaffolded our contigs to generate 30 chromosome-length scaffolds (minimum length 44 Mbp). Then we made two edits based on strong chromosomal block signals from Hi-C alignment and assigned scaffold names based on their respective scaffold sizes (Fig. 1C). This further improved our assembly contiguity by increasing our N50 from 50 Mbp to 125 Mbp (Table 1). Compared to scaffold-level OctDeg1.0 reference, this new assembly increased N50 by more than 10-fold (from 12 Mbp to 125 Mbp) and reduced the number of scaffolds by close to 25-fold (from 7,135 to 288). In the final assembly, 99% of all the genomic material was contained within the 30 largest scaffolds with an average of 6-7 gaps, which reached a chromosome-level assembly. Contamination screening identified a small number of non-target contigs, the removal of which resulted in a final submitted assembly of 3.40 Gb comprising 200 sequences.

Next, we sought to find both the X and Y chromosomes, which are highly repetitive and therefore difficult to assemble. To enable synteny analysis of coding regions, we first built a genome annotation by Oing over the OctDeg1.0 genome annotation to our scaffold assembly. Then, we found the protein-coding region of scaffold1 to have exclusive synteny with the X chromosome of the mouse genome, therefore helping to confidently identify scaffold 1 as chromosome X (Fig. 1D). However, since OctDeg1.0 does not have annotations for Y chromosome genes such as *Sry*, we cannot use the same approach to identify the Y chromosome. Nevertheless, we determined our Y chromosome using three additional lines of evidence. First, by aligning the long and short reads to the masked and unmasked genomes, we found scaffold 1 and 23 to have around half of the coverage for other chromosomes (Fig. 1D).

Next, by overlapping our k-mer set for each chromosome with sequences from a female degu, we found scaffold 23 to have the highest percentage of unique K-mers, suggesting it to be the Y chromosome (Fig. 1E). Finally, we used BLAST program (Camacho et al., 2009) to identify a partial 243-246 bp match between the grey squirrel *Sry* and a region on scaffold 23 (Fig. 1F).

Taken together, these analyses strongly suggest scaffold 23 to be the Y chromosome.

### Repeat and gene annotation

In Fig. 2A, we detailed our genome annotation pipeline used to annotate the repetitive and genetic features of our assembly. This pipeline includes three major steps: 1) Repeat modeling and repeat masking, 2) Transfer-based & *de novo* genome annotation, 3) further enhancement using *de novo* annotation. *De novo* repeat finding by RepeatModeler (Flynn et al., 2020; Tarailo-Graovac & Chen, 2009) generated 1275 consensus repeat sequences. Out of the 1275 consensus repeats, ∼60% (762) could not be classified, while the remaining ∼40% (513) sequences were assigned to 20 transposable element (TE) families (Supplemental Table 2). The largest TE families identified are 141 unknown long terminal repeats (LTRs). Using this *de novo* repeat library, RepeatMasker masked over half (50.07%) of the genome (Table 2). Over 16% of the genome is masked by unclassified repetitive sequences, while retroelements dominate the rest. Among retroelement genomes, 19.31% belong to long interspersed nuclear elements (LINEs), followed by short interspersed nuclear elements (SINEs) at 7.69% and LTRs at 4.16%.

**Table 2.** Interspersed repeat statistics identified from repeatMasker and repeatModeler2. No elements were detected for Penelope, CRE/SLACS, R1/LOA/Jockey, R2/R4/NeSL, RTE/Bov-B, BEL/Pao, Ty1/Copia, En-Spm.

|  | Number of elements | Length occupied | Percentage of sequence |
| --- | --- | --- | --- |
| Retroelements | 4,482,147 | 1,060,590,693 bp | 31.16% |
| SINES | 2,254,854 | 261,862,363 bp | 7.69% |
| LINEs: | 1,229,112 | 657,279,210 bp | 19.31% |
| L2/CR1/Rex | 647 | 64,493 bp | 0.00% |
| L1/CIN4 | 1,228,465 | 657,214,717 bp | 19.31% |
| LTR elements: | 998,181 | 141,449,120 bp | 4.16% |
| Gypsy/DIRS1 | 777 | 508,709 bp | 0.01% |
| Retroviral | 788,195 | 111,011,343 bp | 3.26% |
| DNA transposons | 74,620 | 16,589,778 bp | 0.49% |
| hobo-Activator | 7,655 | 2,267,743 bp | 0.07% |
| Tc1-IS630-Pogo | 57,949 | 10,138,776 bp | 0.30% |
| MULE-MuDR | 165 | 51,784 bp | 0.00% |
| PiggyBac | 8,201 | 3,739,918 bp | 0.11% |
| Unclassified: | 2,218,914 | 567,957,653 bp | 16.69% |
| Total interspersed repeats |  | 1,645,138,124 bp | 48.33% |

**Figure 2.**
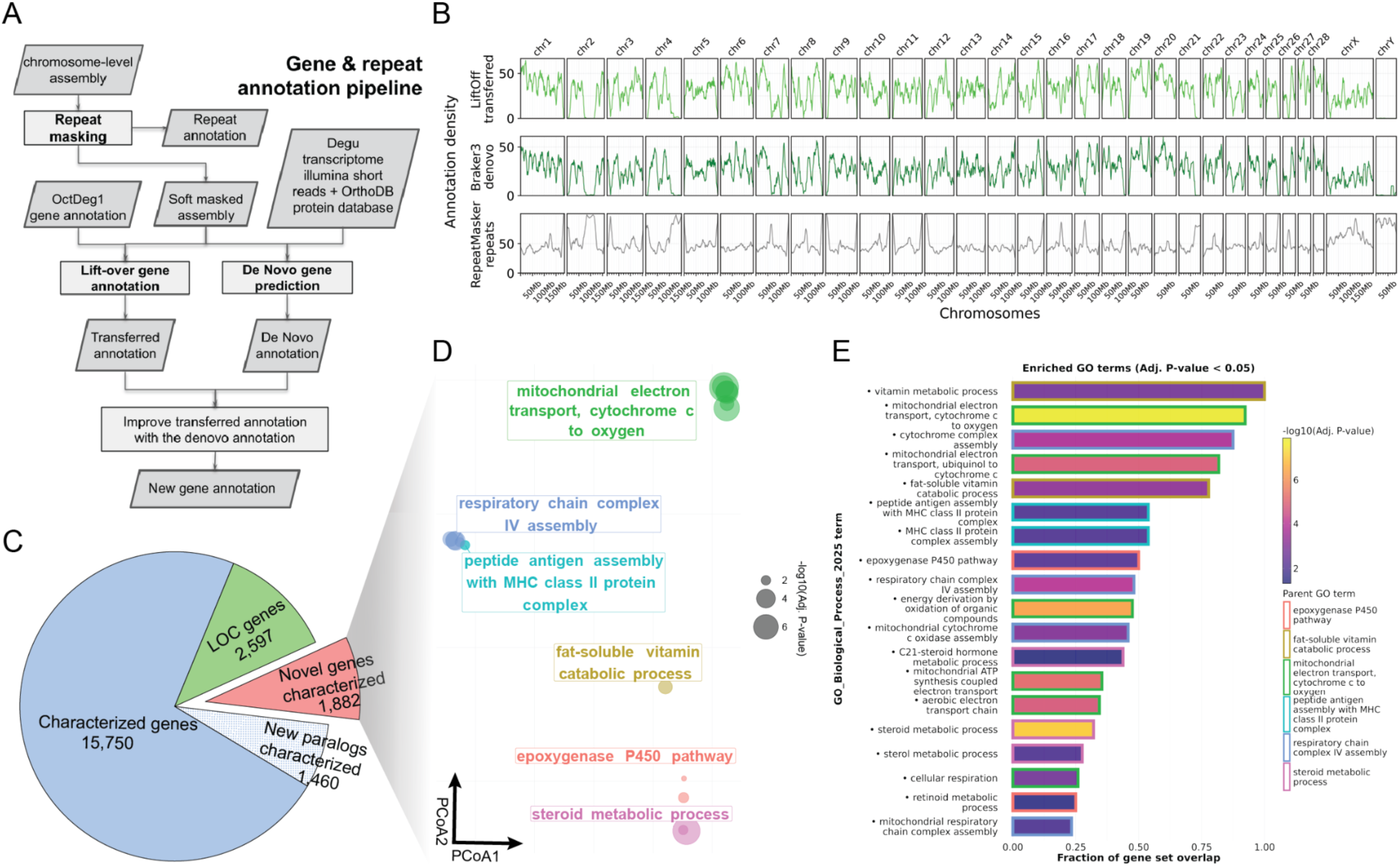
*De novo* genome annotation pipeline improves *Octodon degus* annotation. A) Flowchart of the gene annotation pipeline. B) Whole genome density plot of genes from liftover-based annotation (LiftOff), *de novo* based annotation (Braker3), and repeats from repeatMasker. Pie chart describing our new gene annotation. C) We characterized 1,882 new genes and 1460 new paralogs, which account for 8.68%* and 6.73% of the gene annotation, respectively. D) A scatter plot of the first two PCoA of statistically significant GO terms associated with the newly characterized genes, which are then clustered and labeled by their respective parent GO terms. E) Barplot of the fraction of gene set overlap for each GO term. Filled colors represent adjusted p-values, and border colors are the respective clustered parent GO terms.

Our gene annotation pipeline utilized both liftoff (Shumate & Salzberg, 2021), a liftover-based tool, and Braker3 (Gabriel et al., 2024) , a *de novo* genome annotation pipeline, to build an improved annotation. The liftover-based annotation successfully transferred almost all (99.57% or 25,109 out of 25,217) of non-pseudogene gene features from the OctDeg1.0 annotation to our new genome. Since the liftover-based annotation was quite complete, we utilized the information from the *de novo* annotation to complement the liftover-based annotation and created our final annotation. The *de novo* annotation pipeline successfully identified 26,789 gene features and annotated 23,542 of these features (Supplemental Fig. 5). Interestingly, around 42% of the newly *de novo* annotated gene features have paralogous gene names (Supplemental Fig. 5). The *de novo* annotated gene density is visually congruent with the transferred annotation gene density across the entire genome, suggesting successful *de novo* gene prediction (Fig. 2B). To improve the transfer-based annotation with *de novo* annotation, we built a custom pipeline that added 1882 novel annotated genes and 1460 potential paralogs to the liftover-based annotation. During this process, 40% of genes containing an LOC prefix identifier from the liftover-based annotation were reannotated into functionally interpretable gene symbols (4782 to 2597) (Fig. 2C & Supplemental Fig. 4).

Gene ontology (GO) analysis of the newly characterized genes in the final annotation found enrichment in biological processes involved in mitochondrial electron transport, steroid metabolism, fat catabolic processes, and immune-related pathways (Fig. 2D). Further breakdown of these parent GO terms reveals specific GO pathways involved in peptide antigen assembly with MHC Class II protein complex (Fig. 2E).

<u>Newly assembled regions span repetitive regions and capture additional epigenetic signals</u> To better understand the additional genomic material assembled in OctDeg2.0, we examined the alignment between OctDeg2.0 and OctDeg1.0. Examining the alignments between the two assemblies as a dotplot, we found that all OctDeg2.0 chromosomes have corresponding alignment to OctDeg1.0 (Fig. 3A). The alignment between OctDeg1.0 and OctDeg2.0 spans 2.5 Gbp (2,527,717,165 bp), which accounts for 74.27% of the OctDeg2.0 genome and 83.95% of the OctDeg1.0 genome. Looking at the proportion of alignment across repetitive regions classified by RepeatMasker, we found that most of the alignment falls within the nonrepetitive region of the genome. Specifically, 1.5 Gbp or 61.8% of alignment falls within the nonrepetitive region, and 965 Mbp or 38.2% falls within the repetitive region (Fig. 3B). In addition, we also found that more than 90% of OctDeg2.0 nonrepetitive regions were aligned, as compared to less than 60% of OctDeg2.0 repetitive regions (Fig. 3C). A high alignment percentage in nonrepetitive regions indicates robust congruence between the two assemblies in nonrepetitive regions. Meanwhile, a lower alignment percentage in repetitive regions suggests that OctDeg2.0, as a long-read based assembly, was able to span and recover more repetitive regions.

**Figure 3.**
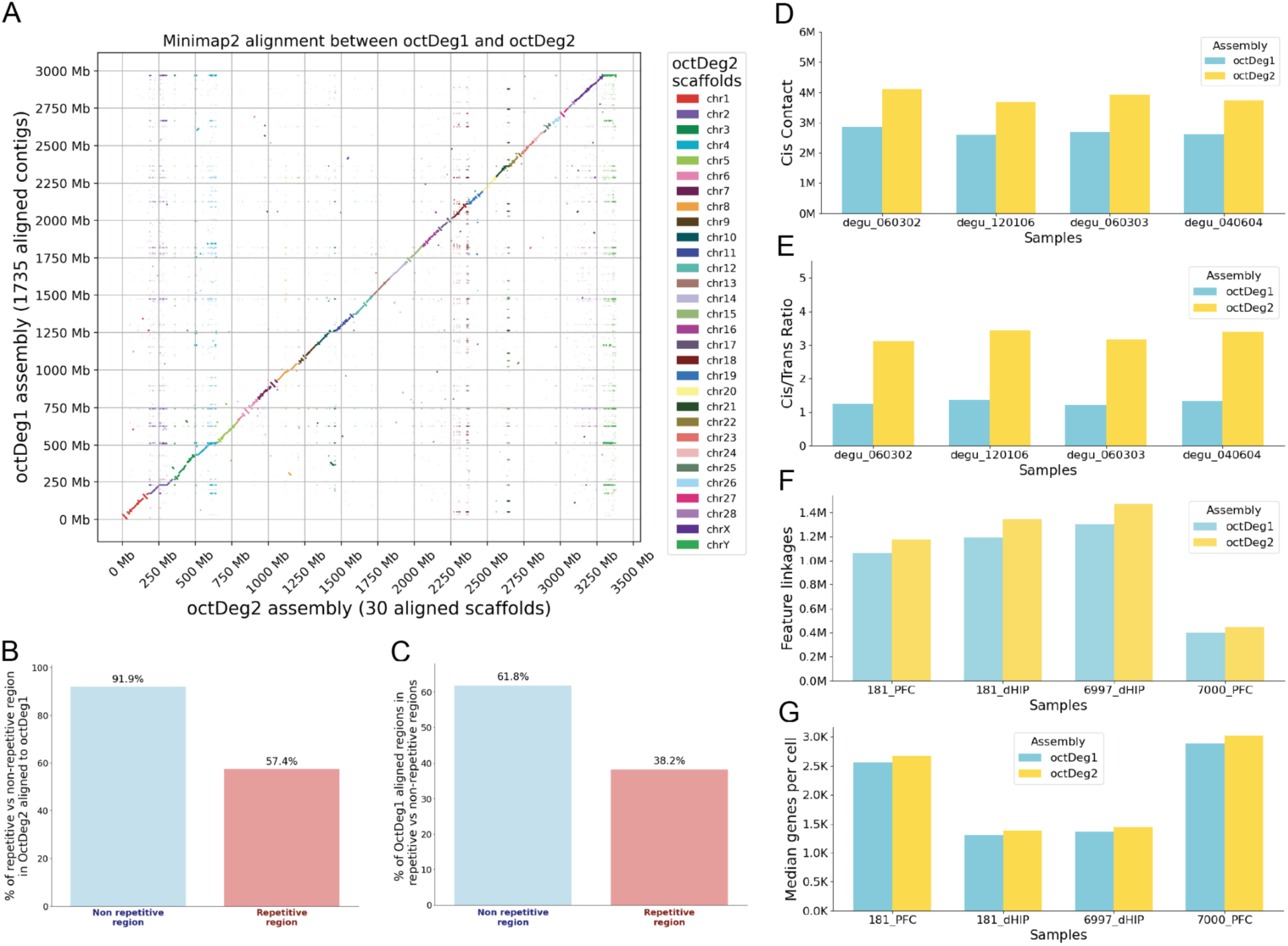
Comparison of genome assemblies reveals the addition of repetitive regions and epigenetic features. A) A scatterplot of the alignment between OctDeg2.0 and OctDeg1.0. The alignment is sorted based on the OctDeg2.0 aligned position and colored by the chromosome aligned. B) A barplot showing the percentage of aligned regions lying within repetitive or nonrepetitive regions of OctDeg2.0. C) A barplot showing the percentage of repetitive and nonrepetitive regions from OctDeg2.0 aligned to OctDeg1.0. Barplots of Cis contact (D) or Cis/Trans ratio (E) from 4 Hi-C samples when aligned to OctDeg1.0 and OctDeg2.0. Barplots of feature linkage (F) or mean genes per cell (G) from 4 Hi-C samples when aligned to OctDeg1.0 and OctDeg2.0.

OctDeg2.0 improved 3D chromosome contact signal and candidate cis-regulatory detection. We analyzed the chromosome contact signal by mapping the Hi-C data from four other degu brain samples to both assemblies. The result revealed a 25% increase in cis contact and more than doubling of the ratio between intra-chromosomal contact over inter-chromosomal contact (cis/trans ratio) in OctDeg2.0 (Fig. 3D & Fig. 3E). This illustrates that OctDeg2.0 enables the capture of more cis-chromosomal contacts, empowering our ability to understand the role chromosome architecture plays in regulating gene expression in degu. Furthermore, we also examined the candidate cis-regulatory element (cCRE) signal by mapping single-cell Multiome ATAC-seq/RNA-seq data (GEO accession: GSE314504) that we generated for the hippocampal tissues from four degu brain samples to both assemblies (Fig. 3F & Fig. 3G). We found consistent increases in the number of feature linkages between accessible regions of the chromosome and gene expression across the 4 degu samples, corresponding to a 10-12% increase when mapped to the OctDeg2.0 genome (Fig. 3F). Thus, OctDeg2.0 enabled us to better identify the links between potential enhancers and their regulatory gene targets. Both improvements in epigenetic mapping illustrate how OctDeg2.0 increases our power to interrogate the role that the non-coding genome plays in gene regulatory programs of the *Octodon degus*.

OctDeg2.0 assembly captured the centromeric regions of the degu genome

We identified 1,315,981 tandem repeats in our genome, a number that lies between those identified in the human reference genome (hg38; 962,790) and the mouse reference genome (GRCm39; 1,637,075). Examining the entire set of tandem repeats by their respective total repeat length and single repeat length, we found a clear enrichment of three single repeat lengths with long tandem repeat lengths at around 195 bp, 349 bp, and 389 bp, which we call tandem repeat stripes (Fig. 5A). Out of the three stripes, 349 bp is especially prominent with multiple ultra-long 349bp monomer tandem repeats with the longest reaching more than 10 Mbp (Fig. 5A), forming a striking tandem repeat stripe pattern. To investigate if these patterns of long tandem repeats in degus are likely centromeric, we examined tandem repeat stripes at known centromeric repeat length in the human hg38 assembly. Indeed, we found two groups of long tandem repeats stripes at 171 and 342 bp, which correspond to the lengths of human centromeric satellite repeats (Supplemental Fig. 9C). To examine whether these ultra-long tandem repeats form higher-order repeats (HORs)—large tandem arrays composed of more basic repeat units characteristic of vertebrate centromeres (Willard, 1991)—we annotated the HOR regions. Unsurprisingly, sites of 349bp tandem repeats and high tandem repeat density regions often correspond with the location of HOR regions (Fig. 4). It is worth noting that many HOR regions also occur at sites of high tandem repeat density, likely corresponding to duplicated HOR repeats at pericentromeres, interspersed HOR repeats at acrocentric short arm, and subtelomeric HOR (Fig. 4). We also observe that many of the tandem repeats and HOR signals are concentrated at contig gaps, suggesting that these repetitive regions are assembly bottlenecks (Fig. 4). These super-long 349 bp tandem repeats are also found in an independently generated degu genome assembly produced using the Verkko assembler, indicating that these ultra-long tandem repeats are unlikely to be assembly artifacts of our pipeline (Supplemental Fig. 14A).

**Figure 4.**
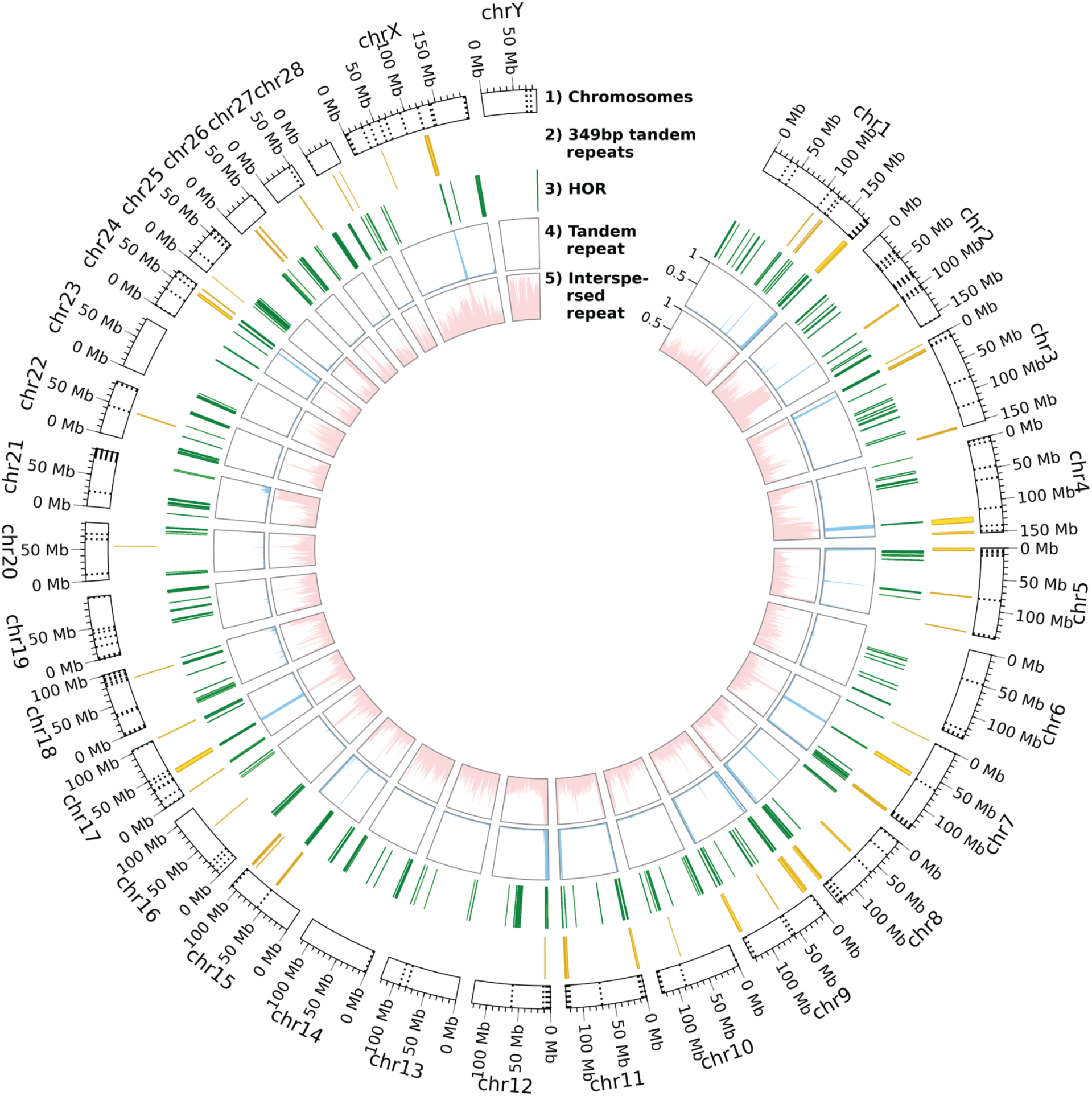
Circosplot of HOR and tandem repeats. Track 1 blocks represent the assigned chromosomes labeled with size and scaffold gaps plotted as dashed lines. Track 2 are the genomic locations of tandem repeats with 349 bp long consensus sequence. Track 3 are the genomic locations of identified high order repeats (HOR). Track 4 is the density plot of the tandem repeats. Track 5 are the density plot of the interspersed repeats.

**Figure 5.**
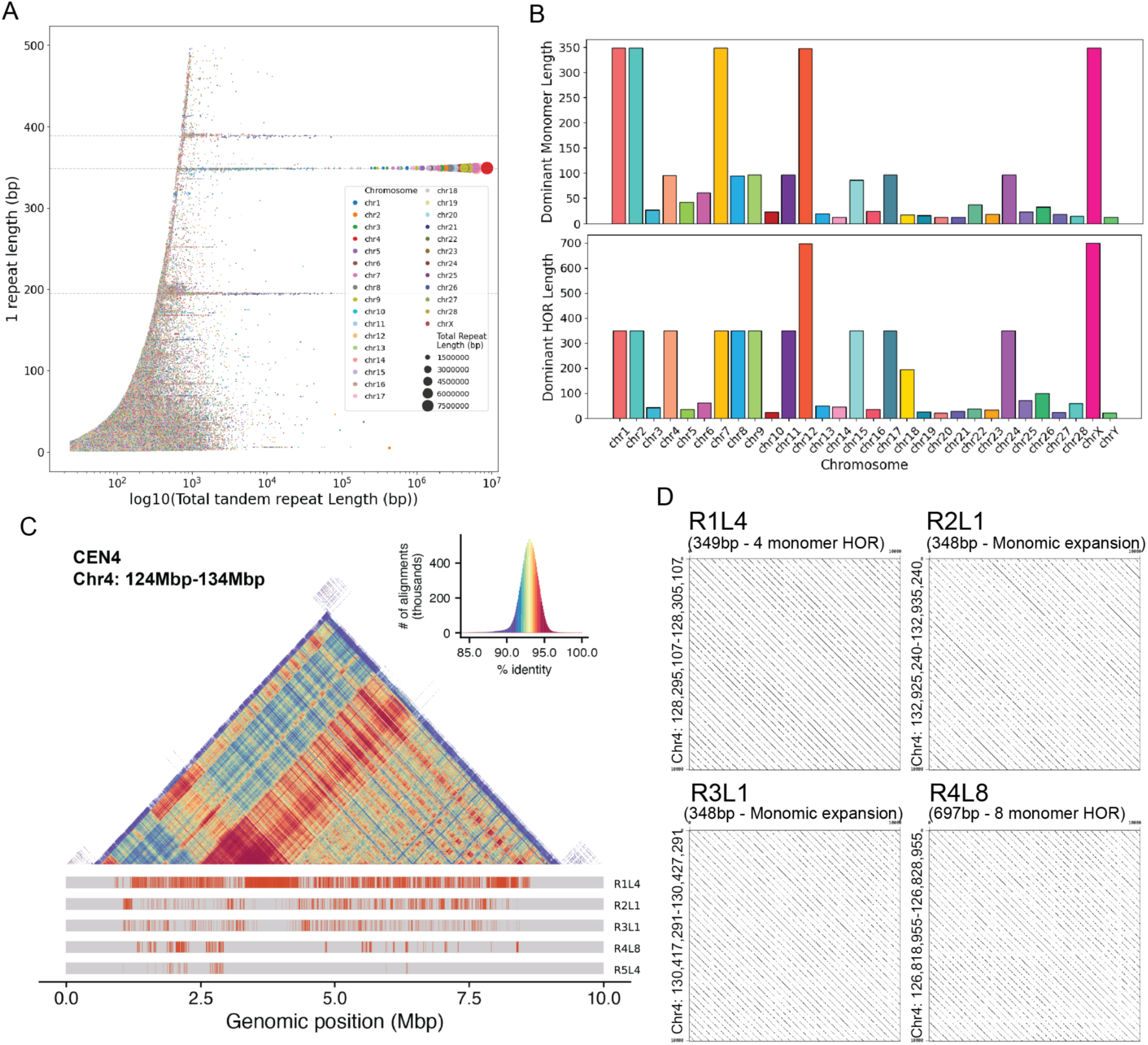
Centromeric repeat analysis reveals HOR of putative centromeric regions. A) A scatter plot of genome-wide tandem repeats plotted by total tandem repeat lengths and monomer lengths and colored by the chromosome origin. B) Barplots of the length of the most abundant monomers (top) and high-order repeats (bottom) for each chromosome. C) Structure and annotation putative CEN4 region. D) Dotplots of repeat patterns from different HOR or monomers found in the putative CEN4 region.

Next, we asked if the tandem repeats with around 349 bp consensus sequence length share more sequence similarity to each other as compared to other tandem repeats in the genome. We compared the 10 longest tandem repeats from each tandem repeat stripes at 349 bp, 389 bp (386 - 390 bp), and 195 bp (193 - 195 bp) in a T-SNE plot (Supplemental Fig. 9A). We find that these 349 bp tandem repeats cluster together in T-SNE plots and not with other tandem repeats, indicating that they share more sequence similarities within each class than with other tandem repeats. We also examined if the 349 bp repeats contained any species-specific characteristic by comparing them to tandem repeats from the genomes of humans, dogs, and other rodents (naked mole rat, guinea pig, mouse, and rat). We used the alignments of the 10 longest tandem repeats from all species to build a sequence similarity tree (Supplemental Fig. 9B). Our analysis reveals that representative tandem repeats from rodents tend to cluster together as opposed to representative tandem repeats from humans and dogs, suggesting some rodent-specific features. Furthermore, motif analysis of these 349bp tandem repeat region reveals an enrichment of binding sites for STAT family, TEAD1, and Zfp809, which are known to be enriched in endogenous retrovirus, LINE / SINE, and other repetitive elements (Supplemental Fig. 9C).

Corresponding to tandem repeat results from tandem repeat finder (Benson, 1999) showing the enrichment of 349 bp repeats, centroAnno (Qi et al., 2025), a centromere-specific annotation program, found 349 bp repeating monomers as most abundant monomers on 5 chromosomes (chr1, chr2, chr7, chrX), and 349 bp repeating HOR as the most abundant HOR on 10 chromosomes (chr1, chr2, chr4, chr7, chr8, chr9, chr11, chr15, chr17, chr24) (Fig. 5b, Supplemental Fig. 7, Supplemental Fig. 8). Interestingly, we find that the 349bp monomers on chr1, chr2, and chr7 remain consistent as the 349bp HOR, whereas the 348bp monomer from chr12 and 349bp monomer from chrX become part of 696bp and 698bp HOR, respectively. This suggests the presence of dimerized satellite repeats on chr12 and chrX. For 7 other unmentioned chromosomes (chr4, chr8, chr9, chr11, chr15, chr17, chr24), the 349 HOR is made up of smaller monomers with lengths of 97-95bp and 86bp. For the 18 chromosomes not mentioned above, their abundant HOR lengths range from 195 to 20 bp (Supplemental Fig. 8). We then specifically examine the self-identity percentage and HOR structure in a 10 mb region of 124 Mbp to 134 Mbp on chr4. We found high percent self-alignment identity, where the self-alignment identity ranges from 90-96% across a 9 Mbp region (Fig. 5C). The region is also characterized by long stretches of three HOR (R1L4, R4L8, R5L4) and two monomeric expansions (R2L1, R3L1) (Fig. 5C). Both the high self-identity and HOR structure suggest this region is a centromeric region spanning 9 Mbp. We then visually confirmed the self-alignment patterns in 10 Kbp regions from 2 HOR and 2 monomers in dotplots. These findings suggested that our assembly was able to recover centromeric regions of the degu genome, with many of them characterized by HOR of length 349bp.

### High levels of inter-chromosomal segmental duplications

Using biser (Išerić et al., 2022), we identified 203,532 segmental duplication (segdup) links covering 453.1 Mbp, corresponding to 13.3% of the OctDeg2.0 assembly. To contextualize this value, we applied the same segdup analysis to the previous assembly (OctDeg1.0), the human genome (GRCh38), the dog genome (canFam6), and eight additional rodent species (Supplemental Table 4). This comparative analysis indicated that the proportion of segmental duplication in OctDeg2.0 falls within the range of other rodent species, which typically spans ∼8–10%, including a closely related guinea pig genome with 9.74% segmental duplication.

We observed a markedly higher number of inter-chromosomal segdup links (165,917) than intra-chromosomal segdup links (37,615) (Supplemental Fig. 12). However, overall, the total genomic span of intra-chromosomal segmental duplication regions (345 Mbp) exceeded that of inter-chromosomal duplications (280.9 Mbp), indicating that intra-chromosomal duplications tend to be longer. A similar pattern was observed in the mouse genome, where inter-chromosomal segdup links (29,362) outnumber intra-chromosomal segdup links (67,612), yet intra-chromosomal regions (222.3 Mbp) span a greater fraction of the genome than inter-chromosomal regions (187.2 Mbp). In contrast, both the guinea pig and human genomes exhibited a higher total span of inter-chromosomal segdup regions relative to intra-chromosomal segdup regions. Consistent with previous studies, the average lengths of intra-chromosomal duplications are greater than that of inter-chromosomal segdup (Supplemental Fig. 12).

Genome-wide visualization of the segdup signal across the whole genome revealed a strong enrichment in the Y chromosome, which exhibited both high segdup density and a strong bias toward inter-chromosomal duplications (Fig. 6 & Supplemental Fig. 11). Specifically, chrY-linked inter-chromosomal segdups connect portions of chr2, chr3, chr4, chr10, and chr26.

**Figure 6.**
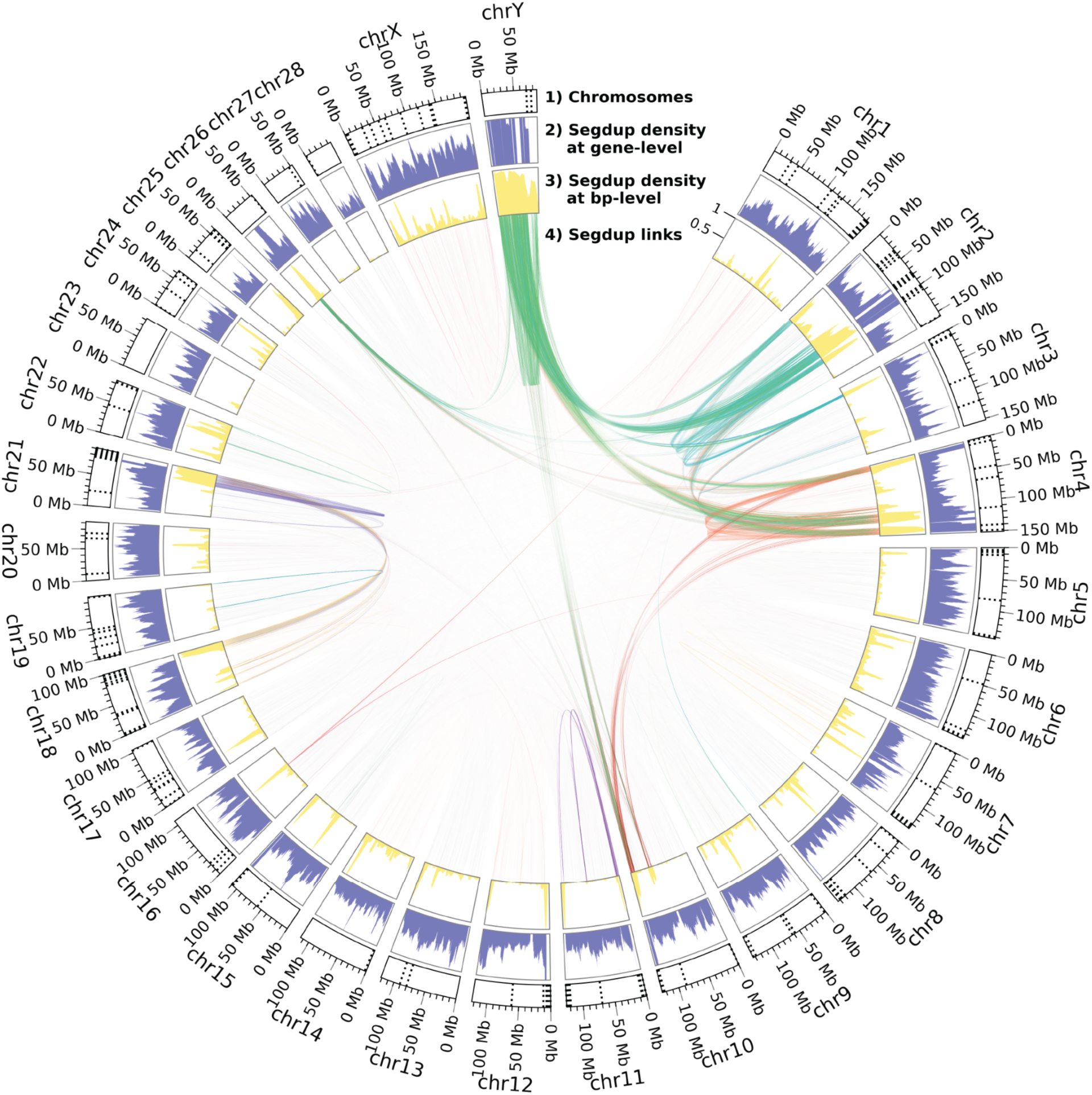
Circosplot of whole-genome segmental duplication (segdup) features reveals extensive Y chromosome-linked inter-chromosomal segdup links. Track 1 blocks represent the assigned chromosomes labeled with size and scaffold gaps plotted as dashed lines. Track 2 contains a segmental duplication density plot calculated as the percentage of segdup overlapping genes in a 5 Mbp rolling window with 100 Kbp steps. Track 3 contains a segmental duplication density plot calculated as the percentage of segdup base pairs in a 5 Mbp rolling window with 100 Kbp steps. Track 4 contains segmental duplication links colored by the chromosome of origins.

Notably, we also find a noticeably inter-chromosomal segdup between chr18 and chr21. To rule out potential assembly artifacts stemming from Hifiasm (Cheng et al., 2021), we reassembled the genome with Verkko (Antipov et al., 2024) and examined its segdup characteristic. Verkko-based assembly revealed a similar pattern of high segmental duplication enriched in the Y chromosome (Supplemental Fig. 12). Interestingly, this mirrors previous findings in humans demonstrating unusually high levels of inter-chromosomal and ampliconic segmental duplications associated with the Y chromosome (Kirsch et al., 2008; Kuroda-Kawaguchi et al., 2001; Skaletsky et al., 2003).

### Segmental duplication pervasive regions contain genes enriched for protein localization to Cajal body

We define segdup pervasive region as the minimum number of segdup regions that make up 80% of the total segdup lengths (Fig. 7A). Using this criterion, we identified the 1,425 longest segdup regions containing 417.9 Mbp of sequences, which includes 1295 gene features equating to 466 unique, non-LOC, genes (Fig. 7A & C). GO enrichment analysis of the 466 genes revealed significant enrichment for pathways related to apoptosis (mitochondrial outer membrane permeabilization; 3/5 genes, padj = 0.041; GO:0097345), nuclear organization (regulation of protein localization to Cajal body; 4/9 genes, padj = 0.020; GO:1904869, GO:1904871), metabolic processes (retinol metabolic process: 5/18 genes, padj = 0.025; GO:0042572; primary alcohol metabolic process: 8/37 genes, padj = 0.005; GO:0034308), and immune responses, particularly interferon signaling and host defense (cellular/response to type II interferon: 8/38 genes, padj ≤ 0.019; GO:0071346, GO:0034341; defense response to virus: 16/161 genes, padj = 0.005; GO:0051607; defense response to bacterium: 14/165 genes, padj = 0.039; GO:0042742). We wondered whether the gene features enriched in the GO terms are functionally relevant and decided to examine the genes enriched in the term Regulation of Protein Localization to Cajal Body pathway. Specifically, the gene features found in this pathway were: Cct3, Cct4-dl1, Cct7-dl1, Larp7-dl6, Larp7-dl7, Larp7-dl2. These genes are directly transferred from OctDeg1.0 annotation (Cct3) or potential “paralogs” added from the *de novo* braker annotation that could be functional paralogs or processed pseudogenes (Cct4-dl1, Cct7-dl1, Larp7-dl6, Larp7-dl7, Larp7-dl2). Looking beyond the segdup pervasive region, we found a total of 21 gene features related to these 4 genes, all of which lies within segdup regions.

**Figure 7.**
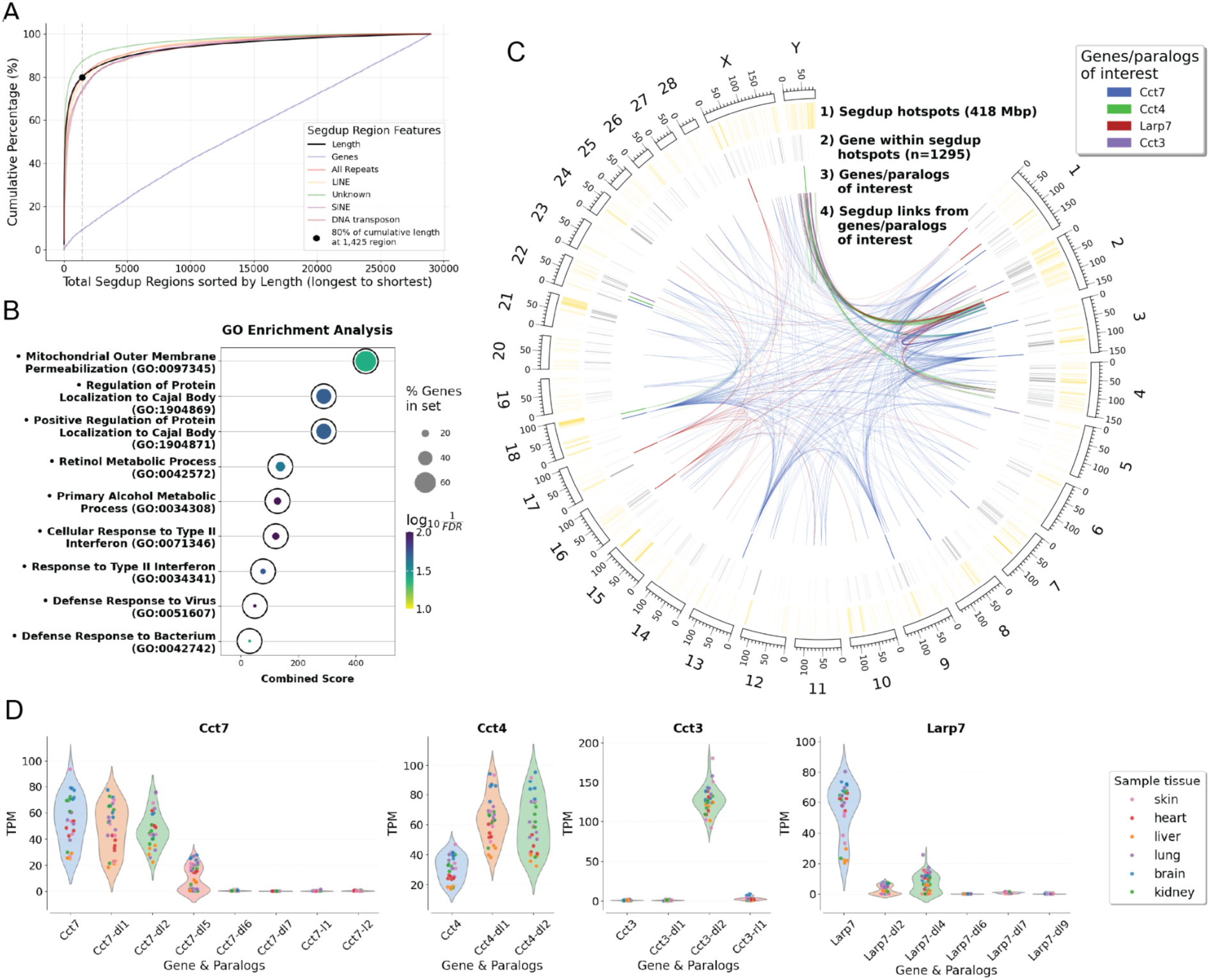
Segmental duplication analysis. A) Percent cumulative plot of lengths from 29,101 merged segdup regions ordered from longest to shortest. We also plotted the percent cumulative plot for other segdup overlapping features: genes, all repeats, LINE repeats, SINE repeats, DNA transposon repeats, and unclassified repeats. The point is the intersect for the 1,425 segdup regions that accounts for 80% of cumulative segdup length, which we define as segdup pervasive region. B) GO enriched biological processes from the 1,295 genes overlapping the segdup pervasive region. C) Circosplot of the location and genes of the segdup pervasive region as well as location and links of four genes/paralogs/pseudogene of interest enriched in the GO term Regulation of Protein Localization to Cajal Body. D) Level of gene expression for four genes/paralogs/pseudogenes of interest in 29 samples across 5 degu organs.

Specifically, we found Cct7, Cct4, Cct3, and Larp7 to have 8, 3, 4, and 6 paralogous genes, respectively. We then visualized all the associated segdup links and positions of these genes (Cct3, Cct4, Cct7, Larp7) and their corresponding gene features (Fig. 7D). We mapped degu transcriptomic data from 29 tissues across five organs (NCBI BioProject PRJNA788430) to determine whether 21 gene features are transcriptionally active. The results show detectable expression of at least one feature for each of the four genes examined (Fig. 7F). These data suggests that segmentally duplicated genes in the degu genome are enriched for functionally active genes involved in Cajal body function.

## Discussion

Here, we present the first chromosome-level genome assembly of *Octodon degus* with both sex chromosomes resolved. Compared to the previous assembly (OctDeg1.0), the new degu reference (OctDeg2.0) exhibits an approximately ten-fold improvement in genome contiguity and shows increased completeness and base-level accuracy relative to a Verkko-based assembly (Supplementary Table 2). Although the primary assembly size exceeds independent genome size estimates, this inflation is consistent with partial haplotype separation by hifiasm and primarily reflects expanded representation of highly repetitive regions.

Importantly, these newly resolved regions substantially improve downstream functional analyses, including chromatin contact maps and the detection of putative regulatory elements.

Although the current assembly demonstrates high contiguity, completeness, and accuracy, additional post-assembly refinement steps could further improve assembly quality. For example, haplotypic duplicate purging, additional rounds of polishing, and post-assembly contamination filtering were not applied in this study. These procedures have been shown to reduce redundancy, correct residual base-level errors, and remove non-target sequences, and their application may further enhance the quality of future *Octodon degus* genome releases.

In addition to improved assembly contiguity, we developed a custom annotation strategy tailored to non-model organisms, for which gene annotations are frequently dominated by poorly characterized loci labeled as “LOC” genes. By integrating homology-based transfer with de novo gene prediction using BRAKER3, our pipeline markedly increases the number of biologically interpretable genes in the degu genome. This approach provides a generalizable framework for improving genome annotations in non-model species and enhances the utility of the degu genome as a resource for comparative and functional genomics.

Owing to their highly repetitive nature, complete resolution of centromeric regions typically requires ultra-long sequencing reads on the order of ∼100 kb. As expected for an assembly based on long but not ultra-long reads (PacBio HiFi, ∼10–20 kb), OctDeg2.0 retains gaps that are predominantly localized to highly repetitive regions and chromosome termini (Fig. 4). Notably, these gaps frequently coincide with regions enriched for higher-order repeat (HOR) structures and extensive tandem repeat arrays, consistent with centromeric and telomeric sequence organization.

Despite these remaining challenges, OctDeg2.0 captures substantial centromeric sequence content and reveals prominent features of degu centromere architecture. We identify exceptionally long tandem repeat arrays, extending up to ∼10 Mb, composed of ∼349 bp repeat units and organized into HOR structures characteristic of centromeric regions (Fig. 5). Although not fully complete, the improved contiguity of OctDeg2.0 enables systematic investigation of centromeric and other highly repetitive regions in the degu genome, providing a foundation for future telomere-to-telomere–level assemblies.

Regions enriched for extensive inter-chromosomal segmental duplications are likely to correspond to pericentromeric domains (Supplementary Table 4). Genome-wide analysis of segmental duplication links revealed a pronounced enrichment of inter-chromosomal duplications connecting the Y chromosome to autosomal regions on chromosomes 2 and 4, and to a lesser extent chromosomes 10 and 26 (Fig. 6; Supplementary Fig. 11A). The major Y-linked autosomal regions on chromosomes 2 (70–110 Mb) and 4 (110–145 Mb) are highly repetitive and exhibit elevated transposable element density (Fig. 2B).

Consistent with their repetitive composition, both long- and short-read mapping to a hard-masked assembly showed markedly reduced coverage across these regions, indicating limited read mappability (Supplementary Fig. 3). These regions are further characterized by low protein-coding gene density and preferential localization within B compartments (Supplementary Fig. 6), features that are hallmarks of pericentromeric chromatin in the mouse genome (Wijchers et al., 2015).

Within each of these large segmental duplication–rich regions, we identify narrower intervals in which segmental duplication levels sharply decrease (chr2: 105–110 Mb; chr4: 124– 134 Mb; Fig. 6, Track 3). Strikingly, these dips coincide with strong higher-order repeat (HOR) signals and abundant 349 bp tandem repeats (Fig. 4, Tracks 2–4), consistent with the location of centromeric cores. Together, these observations support the interpretation that the surrounding segmental duplication–rich regions represent pericentromeric domains flanking centromeres.

The extensive inter-chromosomal duplications observed in the new assembly is consistent with prior reports describing enrichment of segmental duplications in human pericentromeric regions (She et al., 2004). Importantly, similar Y-linked inter-chromosomal segmental duplication patterns were also observed in an independent Verkko-based assembly, indicating that these features are unlikely to be assembler-specific artifacts (Supplementary Fig. 13). To further assess potential scaffolding-related artifacts, we compared HapHiC with an alternative Hi-C–based scaffolder, 3D-DNA. Whereas HapHiC successfully generated chromosome-scale scaffolds, 3D-DNA failed to resolve contigs into clean chromosomal blocks, likely reflecting differences in algorithmic performance in highly repetitive pericentromeric regions rather than artifacts introduced by HapHiC (Supplementary Fig. 14).

As a long-lived, medium-sized rodent, *Octodon degus* represents an attractive system for comparative studies of aging and longevity. Prior comparative analyses across rodents have suggested that enhanced telomerase suppression mechanisms primarily co-evolved with increased body size rather than extended lifespan, raising the possibility that long-lived, smaller-bodied species may rely on telomerase-independent strategies to mitigate cancer risk and promote longevity (Seluanov et al., 2007). Indeed, studies of naked mole rats and blind mole rats have identified distinct tumor-suppressive and longevity-associated mechanisms despite differences in body size and lifespan (Gorbunova et al., 2014).

In naked mole rats, cancer resistance has been attributed in part to the production of exceptionally high–molecular-mass hyaluronan (HMM-HA) mediated by unique amino acid substitutions in HAS2, leading to hypersensitive contact inhibition (Tian et al., 2013). Blind mole rats similarly produce high-molecular-mass hyaluronan, although without exhibiting contact inhibition hypersensitivity, and instead express a unique splice variant of heparanase that represses cancer-promoting activity of the canonical enzyme (Gorbunova et al., 2014; Nasser et al., 2009). In O. degus, genes encoding both hyaluronan synthases and heparanase show robust, tissue-specific expression (Supplementary Fig. 15), suggesting that extracellular matrix–associated pathways may also contribute to longevity-associated phenotypes, although whether degus produce HMM-HA or exhibit altered contact inhibition remains to be demonstrated.

Beyond extracellular matrix pathways, our analysis of segmental duplication–pervasive regions revealed enrichment of expressed genes and putative paralogs associated with mitochondrial outer membrane permeabilization (MOMP), a key regulatory step in the intrinsic apoptotic pathway, as well as proteins involved in Cajal body localization (Fig. 7D; Supplementary Fig. 15A). Segmental duplications are known drivers of copy number variation and elevated nucleotide diversity in mammalian genomes (Perry et al., 2006; Vollger et al., 2022), raising the possibility that such regions in the degu genome represent candidate substrates for lineage-specific regulatory or dosage variation.

Notably, genes associated with Cajal body localization, including Cct7, Cct4, Cct3, and Larp7, exhibit multiple expressed paralogs consistent with duplication-driven expansion. The robust expression of these paralogs across tissues suggests potential functional relevance, although their specific roles in Cajal body regulation or aging-related processes remain to be determined. Together, these findings highlight genomic features that may underlie unique aspects of degu biology and establish a foundation for future functional and comparative studies aimed at dissecting mechanisms of longevity and age-associated disease.

Advances in long-read sequencing technologies have enabled the generation of substantially more contiguous and complete genome assemblies by resolving complex repetitive regions that were previously intractable. High-quality reference genomes are essential for accurately interrogating transcriptomic, genomic, and gene regulatory landscapes, particularly in non-model organisms where such resources remain limited. Despite these advances, de novo genome assembly for non-model species is still technically challenging and often lacks standardized, accessible workflows.

Here, we present a streamlined and reproducible de novo genome assembly and annotation pipeline that can be readily adapted to a wide range of organisms. Applying this framework, we generate the first chromosome-level, sex chromosome–resolved genome assembly of *Octodon degus*, a Chilean rodent of growing interest as an unconventional model for aging and neurodegenerative disease. Together, this resource and accompanying methodology establish a foundation for future functional, comparative, and evolutionary studies in O. degus and provide a practical blueprint for advancing genome assembly efforts in other non-model systems.

## Methods

### Degu tissue procurement

Selected degus are immobilized with head/ears exposed for easy access. Ear of degu are secured using tweezers, exposing the area to cut. We then take the least amount of tissue as necessary. Guided by the tweezer, we make a tangential cut on the apical zone of the ear using the surgical scissors. Target tissue size collected is 3-4mm long and no more than 2mm wide. Deposit tissue in eppendorf tubes and store at 4°C (-20°C for long term (>1week) storage).

### PacBio long-read sequencing

High molecular weight DNA was extracted from Degus ear clip tissues using the NEB Monarch HMW DNA extraction kit for tissue (<u>NEB PN T3010</u>). The HiFi SMRTbell templates were constructed using the SMRTbell prep kit 3.0 (Pacific Biosciences PN 102-182-700) and according to the manufacturer’s protocol. The SMRTbell templates were sequenced on the Pacific Biosciences Sequel II and the Revio platforms.

### Hi-C short-read sequencing

Bulk in situ Hi-C was performed on *Octodon degus* brain tissue using a protocol adapted from established in situ Hi-C workflows from the 4DN consortium. Approximately 20 mg of frozen tissue was homogenized on ice in NIMT buffer (0.25M sucrose, 25mM KCl, 5mM MgCl2, 10mM Tris-HCl pH 7.5, 1mM DTT, 1X Protease Inhibitor (Pierce), and 0.1% Triton X-100) using a Dounce homogenizer (10 strokes with loose pestle A followed by 20 strokes with tight pestle B), taking care to avoid bubble introduction. The homogenate was filtered through a 30μm CellTrics filter (Sysmex, 04-0042-2316) into a LoBind tube (Eppendorf, 22431021) and pelleted (1000 rcf, 10 min at 4°C) (Eppendorf, 5920 R).

The pellet was resuspended in 250 μL of PBS + 1X Protease Inhibitor, and nuclei were counted on a hemocytometer with trypan blue. Nuclei were diluted to 1 million nuclei in 1mL PBS and crosslinked with 1% formaldehyde for 10 minutes at room temperature with gentle rotation. Formaldehyde was quenched by adding 2.5M glycine to a final concentration of 0.2M and rotated for 5 min. Nuclei were centrifuged at 2000xg for 5 min at 4 °C. Pellets were washed once with PBS and centrifuged again at 2000xg for 5 min at 4 °C. Supernatant was discarded and crosslinked pellets were snap-frozen in liquid nitrogen and stored at -80°C or immediately used for conditioning.

Pellets were resuspended in 50 μL of 0.5% SDS with water and incubated for 10 min at 62°C. 170 μl of 1.5% Triton X-100 was added to quench the SDS and incubated at 37°C for 15 minutes. 25 μL of 10x NEBuffer 2 was added and mixed gently. 4 μl of MboI restriction enzyme (25U/μl, 100 U in total) was added and mixed gently. Chromatin was digested overnight (∼16 hours) at 37°C in a thermomixer, shaking at 900 RPM. MboI was heat-inactivated at 62 °C for 20 min. Tubes were cooled to room temperature (∼10 minutes). 50 μL of Fill-in master mix (0.3 mM of dCTP, dTTP, dGTP, and Biotin-14-dATP, and 0.8U of Klenow (M0210L) in water) was added and incubated for 1.5 hours at 37°C in a thermomixer, shaking at 900 RPM. Following fill-in, proximity ligation was performed by adding a T4 DNA ligase master mix (664 µL water, 120 µL 10X T4 ligation buffer, 100 µL 10% Triton X-100, 6 µL BSA (20 mg/mL), 10 µL T4 DNA ligase (400 U/µL), and rotated at room temperature for 4 h.

Nuclei were pelleted at 2,500 × g for 5 min at 4 °C, resuspended in 300 µL PK digestion buffer (10 mM Tris-HCl pH 8.0, 0.5 M NaCl, 1% SDS), and treated with 50 µL proteinase K (20 mg/mL). Samples were incubated 30 min at 55 °C, then overnight at 68 °C. DNA was precipitated by adding 600 µL 100% ethanol and 30 µL 3 M sodium acetate pH 5.2, incubating ≥40 min at −80

°C, and centrifuging at max speed for 15 min at 4 °C. Pellets were washed twice with 800 µL cold 70% ethanol, air-dried for 5 min, and resuspended in 130 µL 10 mM Tris-HCl pH 8.0. Samples were sheared to ∼400 bp on a Covaris M220 using 130-µL microTUBEs with the following parameters: Duty cycle: 10%, Power: 50, Cycles/burst: 200, Time: 70 s. The tube was rinsed with 75 µL water, bringing the total volume to 200 µL. Size selection was performed with 0.55X SPRI beads (110 µL) to remove >500 bp fragments; supernatant retained. Additional 40 µL beads (cumulative 0.75× PEG) to bind 300–500 bp fragments. Beads were washed twice with 700 µL 80% ethanol, dried, and eluted in 300 µL 10 mM Tris-HCl pH 8.0.

Streptavidin T1 or C1 beads (150 µL per sample) were washed twice in 400 µL Tween Wash Buffer (5 mM Tris-HCl pH 8.0, 0.5 mM EDTA, 1 M NaCl, 0.05% Tween-20), resuspended in 300 µL 2X Binding Buffer (10 mM Tris-HCl pH 7.5, 1 mM EDTA, 2 M NaCl), and combined with 300 µL DNA for 20 min at RT. Beads were washed twice at 55 °C for 2 min, then once in 100 µL 1× NEB ligase buffer. 100 uL of End Repair (88 µL 1× T4 ligase buffer, 2 µL 25 mM dNTPs, 4 µL T4 DNA polymerase (3000 U/mL), 5 µL T4 PNK (10,000 U/mL), 1 µL Klenow (exo–, 5000 U/mL)) were added and incubated for 30 min at 24°C. 100 µL of A-Tailing buffer (90 µL 1× NEBuffer 2, 5 µL dATP (10 mM), 5 µL Klenow exo) then incubated 30 min at 37°C.

Beads were resuspended in 50 µL 1× Quick ligation buffer, and 2 µL Quick Ligase plus 1–3 µL Illumina TruSeq LT adapters were added. Samples were mixed and incubated 20 min at 24°C. Beads were washed and resuspended in 89 µL 10 mM Tris-HCl pH 8.0. PCR cycle number was determined by qPCR using a 1:1000 dilution of each Hi-C library. Libraries were amplified using KAPA HiFi polymerase with the following cycling program: 95 °C, 5 min, 98 °C, 20 s, 65 °C, 30 s, 72 °C, 1 min (repeat for n cycles), 72 °C, 7 min. PCR products were pooled (200 µL total), purified with 0.75× SPRI beads (150 µL), washed twice with 1 mL 80% ethanol, dried, and eluted in 100 µL 10 mM Tris-HCl pH 8.0. Libraries were quantified by Qubit and assessed on a TapeStation.

### Whole genome Sequencing of Degus

Using the ear tissues from a male degu (male403), we generated 458,978,150 whole-genome paired-end short reads from Illumina NovaSeq 6000 sequencing platform. With the k-mer (k=21) set generated from these short reads, we used GenomeScope (Vurture et al., 2017) to estimate a genome size of 2,788,820,030 bp and a heterozygosity rate of 0.49%. We then generated 11,327,927 PacBio circular consensus sequencing long reads with an average length of 10,792.23 bp and 4,242,407,455 paired-end Hi-C Illumina short reads with an average length of ∼150 bp from the ear tissue of the same animal, which, based on the approximate genome size, corresponds to 43.8x and 456.3x raw coverage, respectively (Supplemental Table 1). We also sequenced an additional 282,660,623 paired end Hi-C illumina short reads (30.4x raw coverage) to be used as fresh input for scaffolding. The all to all chromatin spatial distance captured by Hi-C reads can be utilized for both assembly phasing and scaffolding.

### Genome assembly

The assembly was built using Hifiasm (v0.25.0-r726) in Hi-C integrated mode with aggressive purging (-l3) and k-mer length set to 21 (-k21) with 43.8x coverage of PacBio long reads and 456.3x coverage of Hi-C short reads (Cheng et al., 2021, 2022, 2024). The input Hi-C read was preprocessed with trim_galore (v0.6.10) on default setting. Hifiasm program outputted a pair of primary and alternative assemblies and a pair of haplotype-resolved assemblies.

The resulting Hifiasm assemblies and the OctDeg1.0 reference were evaluated using custom scripts, BUSCO (v5.7.1), and Merqury (v1.3) to assess contiguity, completeness, and base-level accuracy. BUSCO analysis was performed in genome mode using the glires_odb10 database (Manni et al., 2021). Whole-genome short-read datasets (SRR19145623 and SRR19145629) were downloaded from the NCBI Sequence Read Archive using parallel-fastq-dump (v0.6.7). A combined k-mer set was generated from these short reads using meryl (v1.3), and the resulting meryl database was used as input for Merqury with default parameters to estimate k-mer completeness and consensus base accuracy (Rhie et al., 2020).

### Contig scaffolding

Mitochondrial contigs were removed from the primary assembly before scaffolding. To identify the mitochondrial contigs, we queried the contigs in the primary assembly against the publicly available degu mitochondrial DNA. 33 contigs with close to 100% alignment and similar size to the mitochondrial DNA were removed (Fig. 1B).

Scaffolding of primary contig assembly was performed using HapHiC (v1.0.7), a chromosome number-aware scaffolding tool, using 60.8x coverage Hi-C reads (Zeng et al., 2024). Half of the 60.8x coverage Hi-C reads were sampled from the Hi-C reads used in genome assembly and half of the reads were additional reads sequenced from the same sample. As before, Hi-C reads were preprocessed with Trim_galore at default setting. The preprocessed Hi-C reads are aligned with bwa (v0.7.18-r1243-dirty), duplicate marked with Samblaster (v0.1.26), and filtered for primary, mapped, and non-duplicate read pairs with samtools (v1.21), as shown in the HapHiC documentation (Faust & Hall, 2014; Li et al., 2009; Li & Durbin, 2009). The BAM file is further filtered for greater than ≥1 MAPQ and edit distance ≤3. Finally, the filtered BAM file and mitochondrial filtered primary assembly were input to HapHiC with the number of chromosomes expected to be 30 (28 autosomes, and 2 sex chromosomes). The number of chromosomes for the common degu is well established to be 58, so the haploid number is 29, and 30 is set because we are assembling a male genome with the Y chromosome (George & Weir, 1972).

The resulting scaffold was then manually evaluated in JuiceBox (v2.17.00), in which two apparent scaffold mis-categorizations were corrected (Fig. 1C & Supplemental Fig. 2; Durand et al., 2016). Then the assembly is sorted by length. The final scaffolded assembly contained contigs separated by 100 Ns in the most probable order and orientation. Assembly evaluation tools were applied again to assess the scaffolded assembly.

### Sex chromosome identification & contamination filtering

The X chromosome scaffold was identified by performing synteny analysis against the mouse GRCm39 genome with JCVI tools (Tang et al., 2024). The scaffolded primary assembly is first annotated with a liftover based annotation tool, liftOff v1.6.3 (Shumate & Salzberg, 2021), to transfer the OctDeg1.0 gene annotation to the new scaffolded assembly. The annotation allows us to examine orthologs between the mouse and degu within the coding sequences for synteny analysis and visualization. We identified scaffold 1 as the X chromosome, as it shared exclusive synteny with the mouse X chromosome (Fig. 1D).

The Y chromosome scaffold was identified with three techniques. First, we calculated the long-read and short-read coverage for both the unmasked and hard-masked scaffolded assembly. The long read and short read alignments were performed using minimap2 v2.28-r1209 (Li, 2018; Vasimuddin et al., 2019) and bwa-mem2 v2.2.1 (Li, 2018; Vasimuddin et al., 2019) and, respectively. Then coverage is calculated using bedtools (v2.31.1) makewindow and bedtools coverage (Quinlan & Hall, 2010). When looking at coverage across the hard masked assembly, scaffold 23 exhibited less than half the coverage compared to the other scaffolds. However, the difference in coverage between scaffold 23 and other scaffolds was less in the unmasked assembly, likely due to the repetitive nature of scaffold 23 allowing it to capture more repetitive reads. Second, using the DiscoverY package (Rangavittal et al., 2019), we compared the overlap of k-mers between each the k-mer set of each chromosome and the k-mer set from HiFi reads from a female degu. We found scaffold 23 has noticeably lower k-mer proportion shared with the female k-mer set than that of other scaffolds (scaffold 23 63% vs other scaffolds >95% shared) (Fig. 1F). Third, we used blastn v2.14.1+ (Camacho et al., 2009) to blast Sry sequences from different rodent sequences downloaded from NCBI against our genome.

As part of the NCBI genome submission process, the scaffold-level assembly was screened for potential contaminant sequences using NCBI’s contamination screening pipeline. Contigs flagged as non-target contamination were removed to generate the final submitted assembly.

### Repeat analysis and masking

Repetitive regions were identified and masked using RepeatModeler v2.0.6 (Flynn et al., 2020; Tarailo-Graovac & Chen, 2009) and RepeatMasker v4.1.7-p1 (Flynn et al., 2020; Tarailo-Graovac & Chen, 2009). *De novo* transposable element modeling and family identification were performed on the scaffolded assembly with RepeatModeler in LTR structural discovery mode. RepeatMasker then used the *de novo* repeat library generated from RepeatModeler to mask the scaffolded assembly.

### Genome annotation

Genome annotation was performed using liftOff, a liftover-based gene annotation method, and Braker3 v3.0.8, a hybrid *de novo* gene annotation method (Gabriel et al., 2024a). A complete degu mitochondrial chromosome was added back to the assembly before annotation. LiftOff was used to transfer the annotation from OctDeg1.0 to the new assembly to generate a quick liftover-based annotation. *De novo* genome annotation was also performed with the singularity image from Braker3 v3.0.8 (Brůna et al., 2024; Buchfink et al., 2015; Gabriel et al., 2024, 2024; Kim et al., 2019; Kovaka et al., 2019; Pertea & Pertea, 2020; Quinlan & Hall, 2010; Stanke et al., 2006, 2008), which leveraged both publicly available RNA-seq data collected from different degu organs and prepartitioned OrthoDB vertebrate protein database combined with protein sequences of close relatives, to predict gene features within a *de novo* assembly. The gene features were then functionally annotated by querying corresponding protein sequences with blast+ against the vertebrata UniProt database for gene identity with blast+ and with InterProScan (v5.74-105.0; Jones et al., 2014) against the default InterProScan databases. Finally, agat_sp_manage_functional_annotation.pl from AGAT v1.4.1 (Dainat et al., 2025) was used to functionally annotate our braker3 genome annotation file with the UniProt and InterProScan outputs to create a functionally annotated *de novo* genome annotation file.

We then used the annotated *de novo* genome annotation to complement the liftover-based annotation in three defined scenarios: 1) adding high confidence genes annotated in the *de novo* annotation but not found in liftover-based annotation, 2) renaming or 3) replacing unannotated LOC genes in liftover-based using information from *de novo* annotations. In scenario 1, we add a *de novo* annotated genes to the liftOff based annotation if it satisfy these three criterias: 1) the *de novo* gene that have no coding sequence (CDS) overlap with that of liftOff genes, 2) the CDS length per gene is at least 60% of that of homologous gene found in mice (mm39), 3) and has >0 gene expression (Supplemental Fig. 4D). Since there are more than 4782 LOC genes (24%) in the liftOff-based annotation, we sought to rename or replace these gene features in the next two scenarios with the additional annotation from Braker3. In scenario 2, we rename a liftoff-based LOC gene to the annotated gene from *de novo* annotation if the CDS(s) from the LOC gene shows unique congruence to the CDS from the braker3 *de novo* annotated gene. Congruence is defined as having the same strand direction, start position, and end position between CDS being compared (Supplemental Fig. 4E). In scenario 3, we replace multiple liftOff-based genes with one *de novo* annotated gene if the CDS(s) from the one *de novo* annotated gene has congruence with CDS from multiple liftOff-based genes. During this reannotation of the liftOff-based annotation using our *de novo* annotation we found many annotated genes with the same name. These genes with the same name but at a novel locus could indicate that we annotated a potential novel paralogs or pseudogenes or a false positive annotation due to assembly error. Regardless, we keep these putative paralogous genes in the new annotation but add a suffix to the gene name to give every gene locus a unique gene name. Depending on the scenarios in which putative paralogs were added to the new genome, we added specific identifiers to these potential paralogs to ensure the uniqueness of each gene feature. Specifically, if we found potential paralog(s) from scenario 1, we appended “-dlx” (-like genes from *de novo* annotation) to the gene name where x is the number of the paralog(s) found. For example, since there is already Rpl35 gene annotated in the liftoff-based annotation, if there are 2 additional gene loci where we included Rpl35 in the final annotation with scenario 1, then we would annotate the 2 genesis features as Rpl35-dl1 and Rpl35-dl2 respectively. We appended “-lx” (like genes from renaming) and “-rlx” (like genes from replacing) if potential paralogs were found with scenario 2 and 3 respectively. Our pipeline utilized custom python scripts, pyranges, and AGAT to build our final annotation based on the described guidelines.

### GO enrichment of newly annotated genes

Our final gene annotation could be broken down into 4 groups: 1) The largest group is the original genes transferred from octDeg1, 2) uncharacterized gene loci annotated by LOC gene name, 3) newly characterized genes loci with duplicate names, which we call paralogous genes, 4) and newly characterized gene that’s non-paralogous. We generated the GO terms enriched in the newly characterized genes using the enrichr function of gseapy v1.1.9 (Fang et al., 2023) and human GO_Biological_Process_2025 database. Then we generated the broad categorical parent GO terms from the gseapy go terms to help interpret the biological pathways using the calculateSimMatrix function from the rrvgo v1.18.0 package (Sayols, 2023) and <u>org.Hs.eg.db</u> database (Carlson et al., 2019).

### Genome comparison analysis

We compared our new assembly (OctDeg2.0) with OctDeg1.0 by first examining the location of their alignments and the various epigenetic metrics generated when mapped to both assemblies. To generate the alignment between the 2 assemblies in an efficient manner, we aligned OctDeg1.0 to OctDeg2.0 using minimap2 (-x asm5 --cs). The position of the alignment from the output Pairwise mApping Format (paf) file was then visualized in a dotplot colored by the respective OctDeg2.0 chromosome in panel Fig. 1A. We then overlapped the alignment to the merged position of repeats from repeatMasker to identify the number of aligned basepairs that fall inside and outside of repetitive regions.

To compare the difference in epigenetic signals, we examine 3D conformation with chromosome contact frequency and candidate cis-regulatory elements with feature linkage predictions. To examine the chromosome contact, we aligned Hi-C data from four other degu brain samples using bwa-mem2 (-SP5M -T0) to each assembly. Then we used samtools and pairtools (v1.0.3; Open2C et al., 2024) to calculate the cis and trans contact from the alignment.

### Nuclei preparation from frozen brain tissue for Chromium Single Cell Multiome ATAC + Gene Expression (10x Genomics multiome)

Dorsal hippocampus or prefrontal cortex degu tissues were pulverized in liquid nitrogen using a mortar and pestle on dry ice. Around 20mg of pulverized brain tissue was added to an ice-cold dounce homogenizer containing 1mL of chilled NIM-DP-L buffer (0.25M sucrose, 25mM KCl, 5mM MgCl2, 10mM Tris-HCl pH 7.5, 1mM DTT, 1X Protease Inhibitor (Pierce), 1U/μL Recombinant RNase inhibitor (Promega, PAN2515), and 0.1% Triton X-100). Tissue was dounce homogenized with a loose pestle (5-10 strokes) to shear large pieces, followed by a tight pestle (15-25 strokes) until the solution was uniform. Homogenate was filtered through a 30μm CellTrics filter (Sysmex, 04-0042-2316) into a LoBind tube (Eppendorf, 22431021) and pelleted (1000 rcf, 10 min at 4°C) (Eppendorf, 5920 R). The pellet was resuspended in 1mL NIM-DP buffer (0.25M sucrose, 25mM KCl, 5mM MgCl2, 10mM Tris-HCl pH 7.5, 1mM DTT, 1X Protease Inhibitor, 1U/μL Recombinant RNase inhibitor) and pelleted (1000 rcf, 10 min at 4°C). The pellet was resuspended in 400uL Sort Buffer (1mM EDTA, 1U/μL Recombinant RNase inhibitor, 1X Protease Inhibitor, 1% fatty acid-free BSA in PBS) with 2μM 7-AAD (Invitrogen, A1310). 120,000 7-AAD-positive nuclei were sorted (Sony, SH800S) into a LoBind tube containing Collection Buffer (5U/uL Recombinant RNase inhibitor, 1X Protease Inhibitor, 5% fatty acid-free BSA in PBS). Final collection volumes were determined and 5X Permeabilization Buffer (50mM Tris-HCl pH 7.4, 50mM NaCl, 15mM MgCl2, 0.05% Tween-20, 0.05% IGEPAL, 0.005% Digitonin, 5% fatty acid-free BSA in PBS, 5mM DTT, 1U/μL Recombinant RNase inhibitor, 5X Protease Inhibitor) was added to achieve a 1X solution. Nuclei were incubated on ice for 1 minute, then centrifuged (500 rcf, 5min at 4C) in a swinging bucket centrifuge. Supernatant was discarded and 650uL of Wash Buffer (10mM Tris-HCl pH 7.4, 10mM NaCl, 3mM MgCl2, 0.1%.Tween-20, 1% fatty acid-free BSA in PBS, 1mM DTT, 1U/μL Recombinant RNase inhibitor, 1X Protease Inhibitor) was added without disturbing the pellet followed by centrifuging (500 rcf, 5 min at 4°C) in a swinging bucket centrifuge. Supernatant was removed, and the pellet was resuspended in 7uL of 1X Nuclei Buffer (Nuclei Buffer (10x Genomics), 1mM DTT, 1 U/μL Recombinant RNase inhibitor). 1μL was used for counting on a hemocytometer after staining with Trypan Blue (Invitrogen, T10282). 16-20k nuclei were used for tagmentation reaction and 10x Genomics controller loading. Libraries were then generated following the manufacturer’s recommended protocol (https://www.10xgenomics.com/support/single-cell-multiome-atac-plus-gene-expression).

10x multiome ATAC-seq and RNA-seq libraries were paired-end sequenced on a NextSeq 2000 to assess data quality. If data quality was satisfactory, libraries were deep sequenced on a NovaSeq 6000 to a target depth of ∼50,000 reads per cell for each modality. Raw sequencing was processed using cellranger-arc v2.0.2 (10x Genomics), generating snRNA-seq UMI count matrices for intronic and exonic reads mapping in the sense direction of a gene for OctDeg1 or OctDeg2.0 genome assembly and gene annotations. To examine the feature linkage metric, we pulled data from Cellranger-arc (v2.0.2) outputs from 4 samples aligned to both assemblies (*Cell Ranger ARC Algorithm Overview | Official 10x Genomics Support*, n.d.).

### Tandem repeat analysis

We identify tandem repeats in our assembly using tandem repeat finder (trf v4.09.1) with the recommended parameters (2 7 7 80 10 50 500 -d -l 20) (Benson, 1999). We then visualize all detected tandem repeats by their single repeat length and total repeat length to get a sense of the repeat characteristics for the longest tandem repeats (Fig. 5A). With this scatterplot, we find a three enriched single repeat length with long tandem repeat lengths at around 195 bp, 349 bp, and 389 bp.

### Centromeric region analysis

We found multiple mega base pair sized long tandem repeats, which are likely sites of centromeric regions. Therefore, we use centroAnno (v1.0.2; Qi et al., 2025), a centromere structural analysis tool, with default settings to identify monomer and high order repeats (HOR) structures in the assembly. From the centroAnno output, we summarized the most abundant monomer and HOR for each chromosome (Fig. 5B & Supplemental Fig. 7 & Supplemental Fig. 8). Then we identified regions with both ultra long tandem repeats (∼10 Mbp) from trf and HOR signal from centroAnno as high-confidence centromeric regions. We then further examined a 10 Mbp high confidence centromeric region (124 Mbp - 134 Mbp) on chromosome 4. Using the monomers identified from centroAnno as input, we used HiCAT (v1.2.0; Gao et al., 2023) to annotate the HOR and monomeric expansion sites within this 10 Mbp region. We combine the HiCAT annotation (the bottom horizontal bars of orange annotations) with a self-identity heatmap of this region generated from StainedGlass (v0.6; Vollger et al., 2022) (Fig. 5C). Adobe illustrator 2025 was used to combine the HiCAT annotation with the StainedGlass heatmap. Then we visualized 10 Kbp regions of the top 4 annotated HOR and monomeric expansion with self-alignment dotplots using the Gepard (2.1) tool (Fig. 5D).

### Segmental duplication analysis

We used biser v1.4 (Išerić et al., 2022) to identify all regions of segmental duplications (segdup) in our assembly. We first preprocessed the biser output by removing duplicate links. To maximize the ability to capture evolutionary distant segdups, we did not set a filtering cutoff for minimum error rate between calculated segdups. Then, to assess the whole assembly segdup metrics, we merged all overlapping segdup regions and used the total number of basepairs in the merged segdup regions to calculate the percentage of segdups in our assembly. We repeated this analysis for OctDeg1.0 and 12 assemblies from other rodent species, dog, and human (Supplemental Table 4). To calculate the metrics for inter-chromosomal vs intra-chromosomal segdups, we separated the segdup links into inter-chromosomal or intra-chromosomal segdup links. Then we performed the same analysis as before for each type of segdup links by first merging the segdup regions and then calculating the number of segdup links, total length, average and median length of the segdup links. We repeated the above analysis for guinea pig, mouse, and human (Supplemental Fig. 12). To calculate the metrics for each chromosome, we isolated the intra-chromosomal and inter-chromosomal segdup links for each chromosome. For the inter-chromosomal segdup link calculation of each chromosome, it is important to account for inter-chromosomal segdups to and from each chromosome of interest. Then using each set of segdup links, we calculated the number of links, total alignment error, and total merged length (Supplemental Fig. 11).

To identify contiguous segdup regions, we merged the segdup regions and sorted the segdup regions from longest to shortest segdups regions. We saw that 80% of all segdup base pairs lies with the first 1,425 longest segdups, which represents 4.9% of all segdup regions (29,101). We define these regions as segdup pervasive regions. Using the number of various types of repetitive elements and gene features within these segdup regions, we built a cumulative percentage plot of segdup length, repetitive element, and gene features based on the order of segdup length (Fig. 7A). We used gseapy and human_GO_bp_2025 database to perform GO enrichment analysis on the 1295 genes that lay within the 1425 segdup regions with the whole transcriptome as background gene set. We identified four genes features that are enriched in the GO term of interests and pulled out the genes and paralogs of each gene features. We used pyCirclize (v1.10.1) to plot the positions of the segdup pervasive region and segdup pervasive region overlapping genes as well as the positions of the genes of interest and the segdup links that overlap with genes of interest.

To evaluate the functional potential of enriched gene features and their paralogs, we downloaded publicly available transcriptomic data (SRA Run IDs: SRR17216293–SRR17216321) from 29 samples across five tissues for gene expression analysis. We quantified expression of all gene features in each sample using Salmon (v1.10.3; Patro et al., 2017) in mapping-based mode with the whole assembly as decoy (Fig. 7D). TPM values for genes of interest were then plotted in Python.

### 3D chromosome contact analysis

Generated Hi-C were mapped to both assemblies to assess both assemblies’ ability to capture chromosome contact information. The Hi-C reads were first trimmed with trim_galore in paired mode at default setting. Then the reads were mapped to reference genome (OctDeg1.0 and OctDeg2.0) with bwa-mem2. Then the alignments are processed using pairtools parse [-- min-mapq 40 --walks-policy all --max-inter-align-gap 30] and pairtools sort to extract sorted Hi-C pairs. Then we use pairtools dedup [--mark-dups --output-stats] to get deduplicated Hi-C cis and trans contact information.

### AB compartment and DNA methylation analysis

We used the 15-fold Hi-C data used for genome scaffolding for AB compartment calculation. To start off, the 15-fold Hi-C data was trimmed with trim_galore then aligned with bwa-mem2. Then the alignment was name-sorted with sambamba. We then used HiCExplorer v3.7.2 package (Lee et al., 2021; Thorsson et al., 2018; Wolff et al., 2018) to build the Hi-C matrix (hicBuildMatrix), correct the matrix (hicCorrectMatrix), then call A/B compartment (hicPCA) at 100 kb bin size. DNA methylation fraction was calculated from mapped single cell methyl Hi-C data from one sample (6834_PFC) processed using the YAP pipeline.

### Circos plot visualizations

We used circos plots to visualize the genome-wide characteristics of our assembly. Python package pyCirclize v0.3.0 was used to plot the cicosplots in Figure 4, Figure 6, Figure 7C and supplemental Figure 6. The outermost chromosome tracks were plotted with the Circos function. The genomic features in Figure 4 (track 2, 3), Figure 7C (track 1,2,3) and supplemental Figure 6 (track 2, 3, 4, 5, 6) were plotted with the “track.genomic_features” function. To visualize density information in Figure 4 (track 4, 5), Figure 6 (track 2, 3), we first calculate the percentage of bases satisfying the criteria of interest within each rolling 5 Mbp window with a step of 100 Kbp. Then we use the “track.fill_between” function to plot the calculated density. To visualize the GC content (Supplemental 6 track 9), we used samtools to calculate GC content in each rolling 10 Kb window with step of 5 Kb. Then we used fill_between to visualize the GC content centered around the average genome-wide GC content. We also use the fill_between function to visualize the AB compartment (Supplemental 6 track 7) and CG methylation fraction data (Supplemental 6 track 8). The CG methylation frac track (Supplemental Fig. 6 track 7) was plotted using the data from the bigwig file for each 10 kb window. For the segdup links in Figure 6 (track 4) and Figure 7C (track 4), we use the “circos.link” function to plot the information from bedpe files.

## Data Availability

Data produced in this study are linked to Bioproject PRJNA1393203 with Biosample SAMN54286841. Long read sequences have been submitted to the SRA database. Genome assembly have been deposited at Genbank under the accession JBTJBO000000000. Functional data has been submitted to NCBI GEO under accession number GSE314504 for 10x Single Cell Multiome data and GSE314503 for Hi-C data.

## Code Availability

Scripts used for data processing and analysis are available on github: https://github.com/JHCCoder/genome-building-pipeline

## Acknowledgement

We thank Yuwu Chen for computational support on TSCC at the San Diego Supercomputer Center (SDSC), Dr. Zhenmiao Zhang for helpful discussions and advice on this work, and Dr. Ting Wang for facilitating the collaboration with Dr. Juan Macias, Dr. Chad Tomlinson.

